# Crystal Structure of Histone Deacetylase 6 Complexed with (*R*)-Lipoic Acid, an Essential Cofactor in Central Carbon Metabolism

**DOI:** 10.1101/2023.08.08.552419

**Authors:** Paris R. Watson, Juana Goulart Stollmaier, David W. Christianson

## Abstract

The enzyme cofactor (*R*)-lipoic acid plays a critical role in central carbon metabolism due to its catalytic function in the generation of acetyl-CoA, which links glycolysis with the tricarboxylic acid cycle. This cofactor is also essential for the generation of succinyl CoA within the tricarboxylic acid cycle. However, the biological functions of (*R*)-lipoic acid extend beyond metabolism owing to its facile redox chemistry. Most recently, the reduced form of (*R*)-lipoic acid, (*R*)-dihydrolipoic acid, has been shown to inhibit histone deacetylases (HDACs) with selectivity for the inhibition of HDAC6. Here, we report the 2.4 Å-resolution X-ray crystal structure of the HDAC6–(*R*)-dihydrolipoic acid complex, and we report a dissociation constant (K_D_) of 350 nM for this complex as determined by isothermal titration calorimetry. The crystal structure illuminates key affinity determinants in the enzyme active site, including thiolate-Zn^2+^ coordination and S-π interactions in the F583-F643 aromatic crevice. This study provides the first visualization of the connection between HDAC function and the biological response to oxidative stress: the dithiol moiety of (*R*)-dihydrolipoic acid can serve as a redox-regulated pharmacophore capable of simultaneously targeting the catalytic Zn^2+^ ion and the aromatic crevice in the active site of HDAC6.

---

The well-known enzyme cofactor (*R*)-lipoic acid ((6*R*)-6,8-dithiooctanoic acid) plays a critical role in central carbon metabolism due to the facile redox chemistry of its disulfide moiety (Figure 1).^1-3^ A prime example includes catalysis by the pyruvate dehydrogenase complex that links glycolysis with the tricarboxylic acid (TCA) cycle.^4-7^ (*R*)-Lipoic acid is covalently tethered to a flexible lysine residue in the E2 subunit of pyruvate dehydrogenase, and in this context is referred to as (*R*)-lipoamide. When bound to the E1 subunit, (*R*)-lipoamide is acetylated to yield the reduced form of the cofactor, acetyl-(*R*)-dihydrolipoamide, which then swings back to the E2 subunit where it acetylates Coenzyme A (CoA) to yield acetyl-CoA and free (*R*)-dihydrolipoamide. Acetyl-CoA then enters the tricarboxylic acid (TCA) cycle and (*R*)-dihydrolipoamide is oxidized in the E3 subunit to regenerate (*R*)-lipoamide. The same reaction sequence is catalyzed by the α-ketoglutarate dehydrogenase complex in the TCA cycle, where E2-linked (*R*)-lipoamide is required for the generation of succinyl-CoA.^8,9^

**Figure 1.**
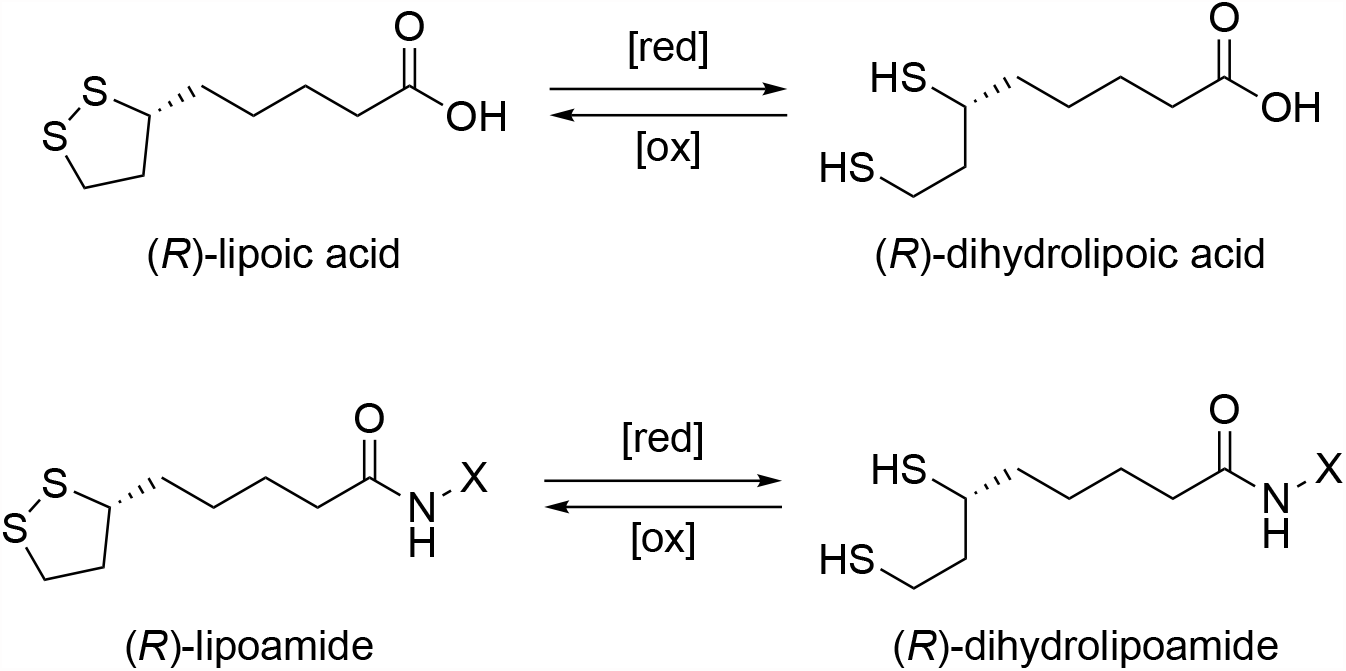
Molecular structure of (*R*)-lipoic acid. The oxidized and reduced states of (*R*)-lipoic acid are shown (top); [red] = reduction, [ox] = oxidation. When attached to the side chain of a lysine residue (X) in the E2 subunit of pyruvate dehydrogenase, the cofactor is referred to as (*R*)-lipoamide in the oxidized state and (*R*)-dihydrolipoamide in the reduced state (bottom); the cofactor is also referred to as (*R*)-lipoamide when amidated to form a simple carboxamide (X = H). The C8 thiol group of (*R*)-dihydrolipoamide undergoes reversible acetylation in the biosynthesis of acetyl-CoA. Oxidation of (*R*)-dihydrolipoamide in the E3 subunit of pyruvate dehydrogenase regenerates the disulfide linkage of (*R*)-lipoamide for another catalytic turnover.

Beyond central carbon metabolism, free (*R*)-lipoic acid (or the racemic mixture of free (*R*)- and (*S*)-lipoic acid, designated “(*R/S*)”) is involved in diverse biological processes. It is an antioxidant that can be used to manage oxidative stress in chronic human disease, such as diabetic neuropathy.^10^ When used in this manner, (*R*)-lipoic acid has been shown to lower blood triglyceride concentrations, although this effect has not been universally observed.^11-13^ Due to its neuroprotective antioxidant effects, (*R*)-lipoic acid is considered as a possible nutraceutical for Alzheimer’s disease therapy;^14^ moreover, (*R*)-lipoic acid is efficacious in animal models of neurodegeneration as well as other oxidative stress-related disorders such as ischemia-reperfusion injury and cataract formation.^15,16^

In a 2023 *Nature Communications* paper, Lechner and colleagues report that (*R*)-dihydrolipoic acid is an inhibitor of the Zn^2+^-dependent histone deacetylases (HDACs), with 5– 15-fold selectivity for inhibition of HDAC6 (EC_50_ = 1 μM).^17^ Neither the oxidized form of the inhibitor, (*R*)-lipoic acid, nor its stereoisomer (*S*)-dihydrolipoic acid, are effective inhibitors. HDAC6 is a class IIb enzyme^18^ predominantly localized in the cell cytosol,^19-22^ where it serves as a tubulin deacetylase and Tau deacetylase.^23,24^ The cell cytosol is a reducing environment, so the disulfide linkage of (*R*)-lipoic acid will exist here predominantly in the reduced form as (*R*)-dihydrolipoic acid with C6 and C8 thiol groups.^25^ Accordingly, Lechner and colleagues hypothesize that one or both of the thiol groups coordinate to the catalytic Zn^2+^ ion in the active site of HDAC6 (as well as other HDAC isozymes).^17^ In the peer review file associated with this paper, the authors state that they were unable to crystallize any HDAC–(*R*)-dihydrolipoic acid complex to conclusively establish the structural basis of inhibition. Of note, contrasting proposals suggest that the carboxylate group of (*R*)-lipoic acid serves as the Zn^2+^-binding moiety.^26,27^ Ambiguities regarding the inhibitory binding mode of (*R*)-dihydrolipoic acid must be resolved to fully understand the biological activity of this critical enzyme cofactor.

Here, we resolve these ambiguities by reporting the 2.4 Å resolution X-ray crystal structure of the HDAC6–(*R*)-dihydrolipoic acid complex. We also report the dissociation constants (K_D_) of (*R/S*)-dihydrolipoic acid, (*R*)-lipoic acid, (*R/S*)-dihydrolipoamide (X = H in Figure 1), and (*S*)-dihydrolipoic acid as measured using isothermal titration calorimetry. The naturally occurring (*R*)-dihydrolipoic acid stereoisomer exhibits the tightest binding with K_D_ = 350 nM, and the crystal structure of the HDAC6 complex reveals that C8–S^−^•••Zn^2+^ coordination and C6-SH••• aromatic interactions are the primary affinity determinants in the active site.

## Results

### Crystal Structure

The structure of the HDAC6–(*R*)-lipoic acid complex was determined at 2.4 Å resolution in space group *P*2_1_2_1_2_1_ with two independent monomers A and B in the asymmetric unit. The binding of (*R*)-dihydrolipoic acid to HDAC6 does not trigger any significant structural changes in either monomer and the root-mean-square deviation of 325 Cα atoms is 0.189 between monomer A in the inhibitor-bound and unliganded (PDB 5EEM)^21^ enzyme structures. Electron density maps show that the inhibitor binds in the active site with only the C8 thiol group coordinated to the catalytic Zn^2+^ ion (average S•••Zn^2+^ distance = 2.35 Å) (Figure 2). Coordination of a thiol group to Zn^2+^ lowers the pKa of the thiol group from ∼8.5 to ∼6 and thereby facilitates ionization,^28^ yielding a potent thiolate-Zn^2+^ charge-charge interaction. The Zn^2+^-bound thiolate is also stabilized by a hydrogen bond with catalytic tyrosine Y745.

**Figure 2.**
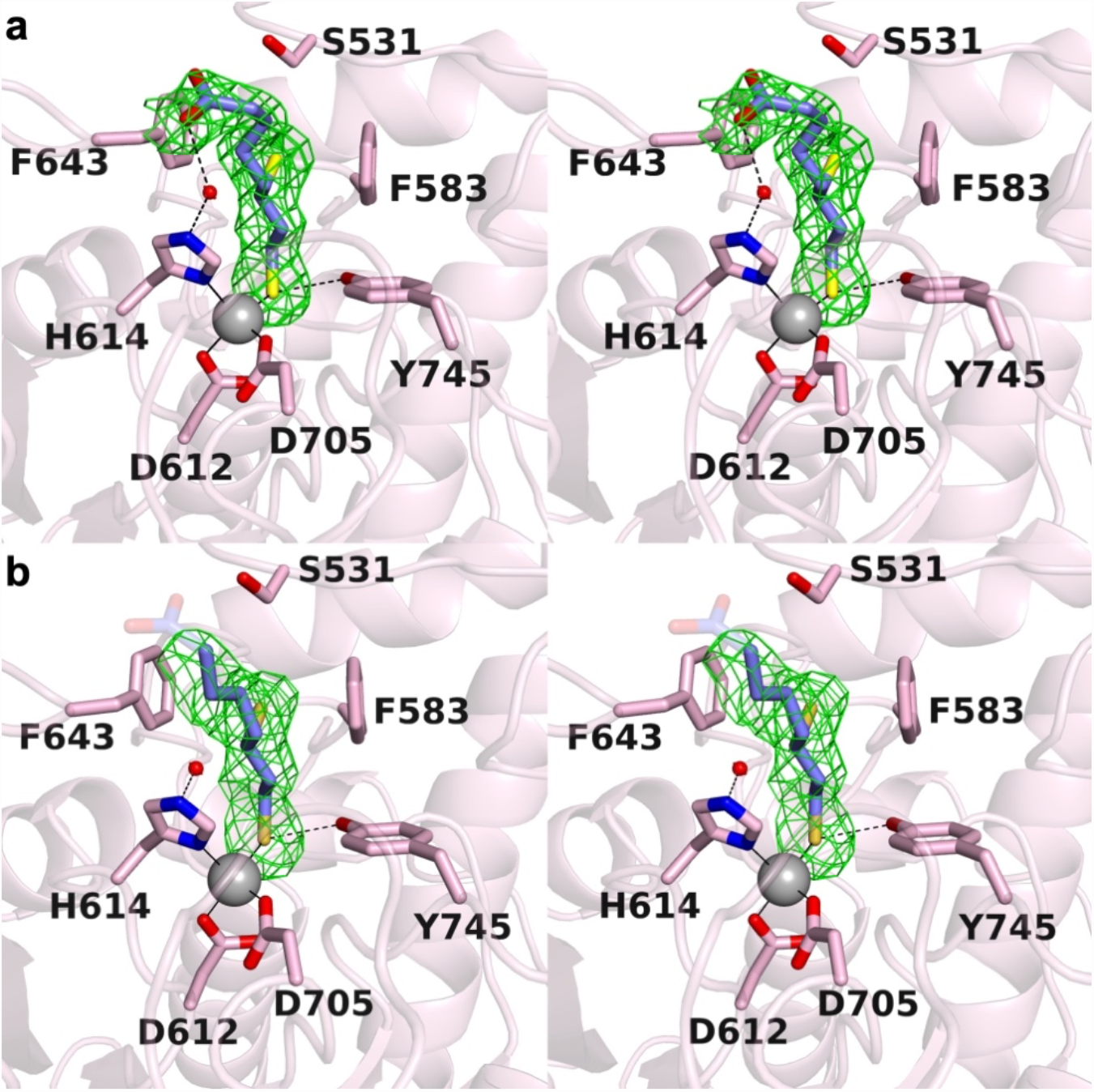
Stereoviews of electron density maps showing (*R*)-lipoic acid bound in the HDAC6 active site. Atomic color codes are as follows: C = mauve (HDAC6) or blue ((*R*)-lipoic acid), N = blue, O = red, Zn^2+^ = gray sphere; metal coordination and hydrogen bond interactions are indicated by solid and dashed black lines, respectively. (a) Polder omit map contoured at 3.2s showing (*R*)-lipoic acid bound in the active site of monomer A. (b) Polder omit map contoured at 3.2s showing (*R*)-lipoic acid bound in the active site of monomer B. Here, the terminal carboxylate group is disordered and shown as a “ghost” image for reference; these atoms are not included in the final model.

The C6 thiol group of (*R*)-dihydrolipoic acid resides in an aromatic cleft formed by F583 and F643, where it engages in S-π interactions.^29,30^ The C6 sulfur atom is located 3.7 Å and 4.0 Å from the aromatic ring centroids of F583 and F643, respectively, in monomer A. This is the first example of an S-π interaction in the aromatic crevice of HDAC6, where more usually the aromatic linker groups of inhibitors typically pack.^21,31-33^ Finally, the carboxylic acid moiety of (*R*)-dihydrolipoic acid is ionized as the negatively charged carboxylate under physiological conditions and extends out of the active site. In monomer A, the carboxylate group forms a hydrogen bond with a water molecule that, in turn, forms a hydrogen bond with Zn^2+^ ligand H614 (Figure 2a). The carboxylate group is disordered in monomer B (Figure 2b). Clearly, the carboxylate group does not engage in direct Zn^2+^ coordination in either monomer, though the carboxylate-H_2_O-H614 hydrogen bond network in monomer A might be viewed as an indirect interaction with Zn^2+^.

### Isothermal titration calorimetry (ITC)

To gain further insight into the binding affinity of lipoic acid derivatives to HDAC6, we studied the thermodynamics of enzyme-inhibitor association using ITC. The dissociation constants (K_D_) determined for each enzyme–inhibitor pair (Figure 3) follow a similar trend to the reported EC_50_ values.^17^ The preferred stereoisomer, (*R*)-lipoic acid, binds with a K_D_ value approximately one-half that of the K_D_ values measured for racemic (*R/S*)-lipoic acid and (*R/S*)-lipoamide. This is consistent with the binding of only the (*R*)-stereoisomer from the racemic mixture; binding was not detected for the (*S*)-stereoisomer.

**Figure 3.**
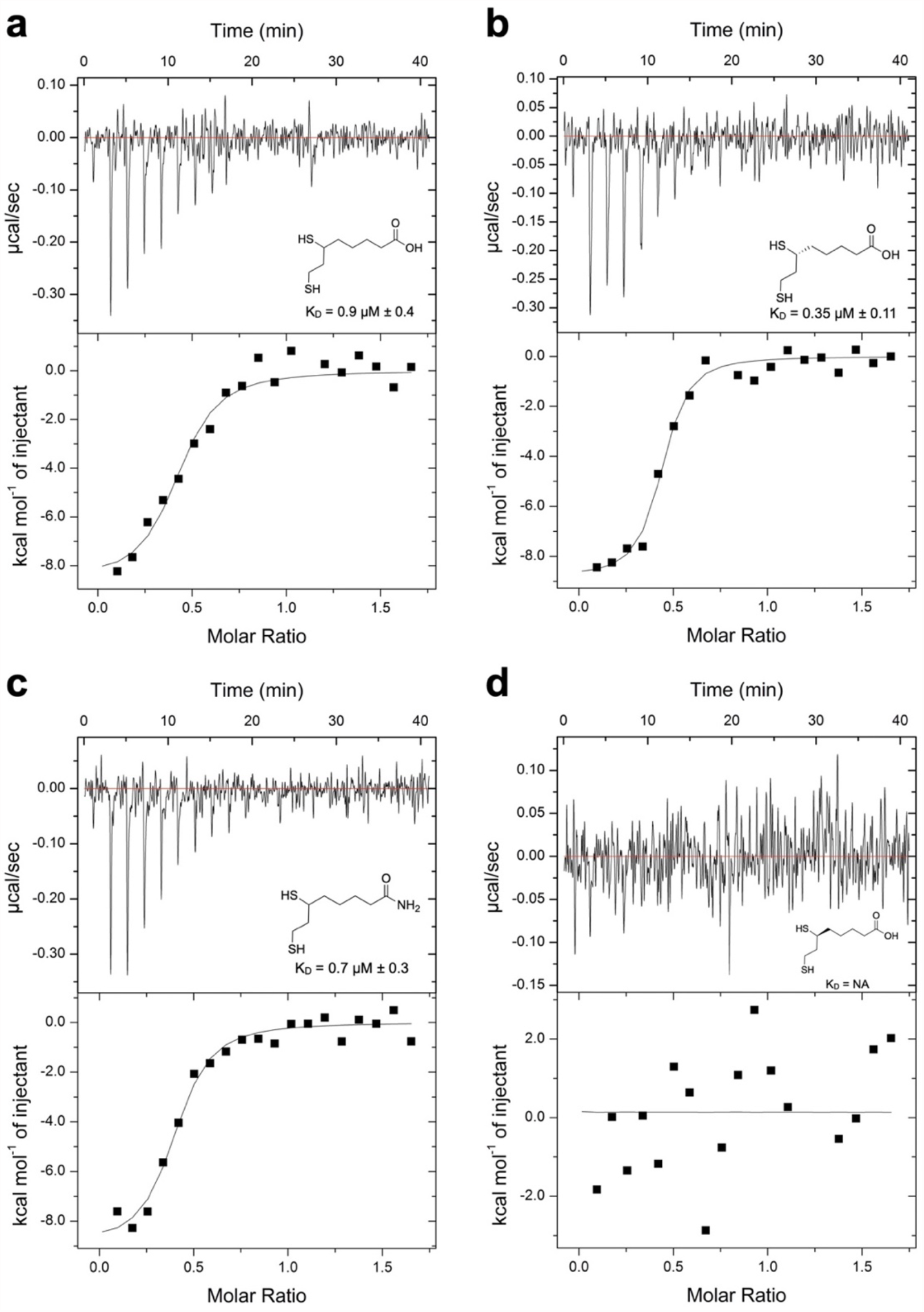
Isothermal titration calorimetry. Enthalpograms for the binding of (a) (*R/S*)-lipoic acid, (b) (*R*)-lipoic acid, (c) (*R/S*)-lipoamide, and (d) (*S*)-lipoic acid to HDAC6.

## Discussion

Perhaps the most prominent feature in the crystal structure of the HDAC6–(*R*)-lipoic acid complex is the coordination of the inhibitor C8 thiolate group to the catalytic Zn^2+^ ion. A handful of crystal structures have been determined of deacetylases complexed with inhibitors bearing thiol Zn^2+^-binding groups, such as the HDAC8–Largazole complex^34,35^ and HDAC6 complexes with thiol-containing macrocyclic octapeptides.^36,37^ Upon coordination to Zn^2+^, the pKa of the thiol group is lowered so as to facilitate ionization, thereby yielding a potent thiolate-Zn^2+^ charge-charge interaction. The ideal geometric parameters for thiolate-Zn^2+^ coordination were first outlined by Chakrabarti through analysis of cysteine-metal ion coordination interactions in the Protein Data Bank:^38^ the average S^−^•••Zn^2+^ separation is 2.1 Å, the average C–S^−^•••Zn^2+^ angle is 108°, and the preferred C–C–S^−^•••Zn^2+^ dihedral angles are 180° and ± 90°. In monomers A and B of the HDAC6–(*R*)-lipoic acid complex, the S^−^•••Zn^2+^ separations are 2.3 Å and 2.4 Å, the C– S^−^•••Zn^2+^ angles are 117° and 122°, and the C–C–S^−^•••Zn^2+^ dihedral angles are 56° and 37°, respectively. While these parameters deviate from the ideal parameters outlined above, they nonetheless allow for high affinity binding.

A second prominent structural feature of the HDAC6–(*R*)-dihydrolipoic acid complex involves the C6 thiol group, which engages in S-π interactions in the F583-F643 aromatic crevice. There are three possible orientations for energetically-favorable S-π interactions (Figure 4a),^37^ two of which are thiol interactions with the face of the aromatic ring. The C6 thiol group of (*R*)-dihydrolipoic acid clearly interacts with the faces of the F583 and F643 aromatic rings (Figure 4b). The geometries of thiol-aromatic interactions with F583 and F643 are generally consistent with energetically-favorable interactions observed between the thiol side chain of cysteine and aromatic residues in an analysis of protein structures contained in the PDB, as well as molecular orbital calculations showing that the positioning of an alkylthiol moiety above an aromatic ring centroid is most preferable.^28,38^

**Figure 4.**
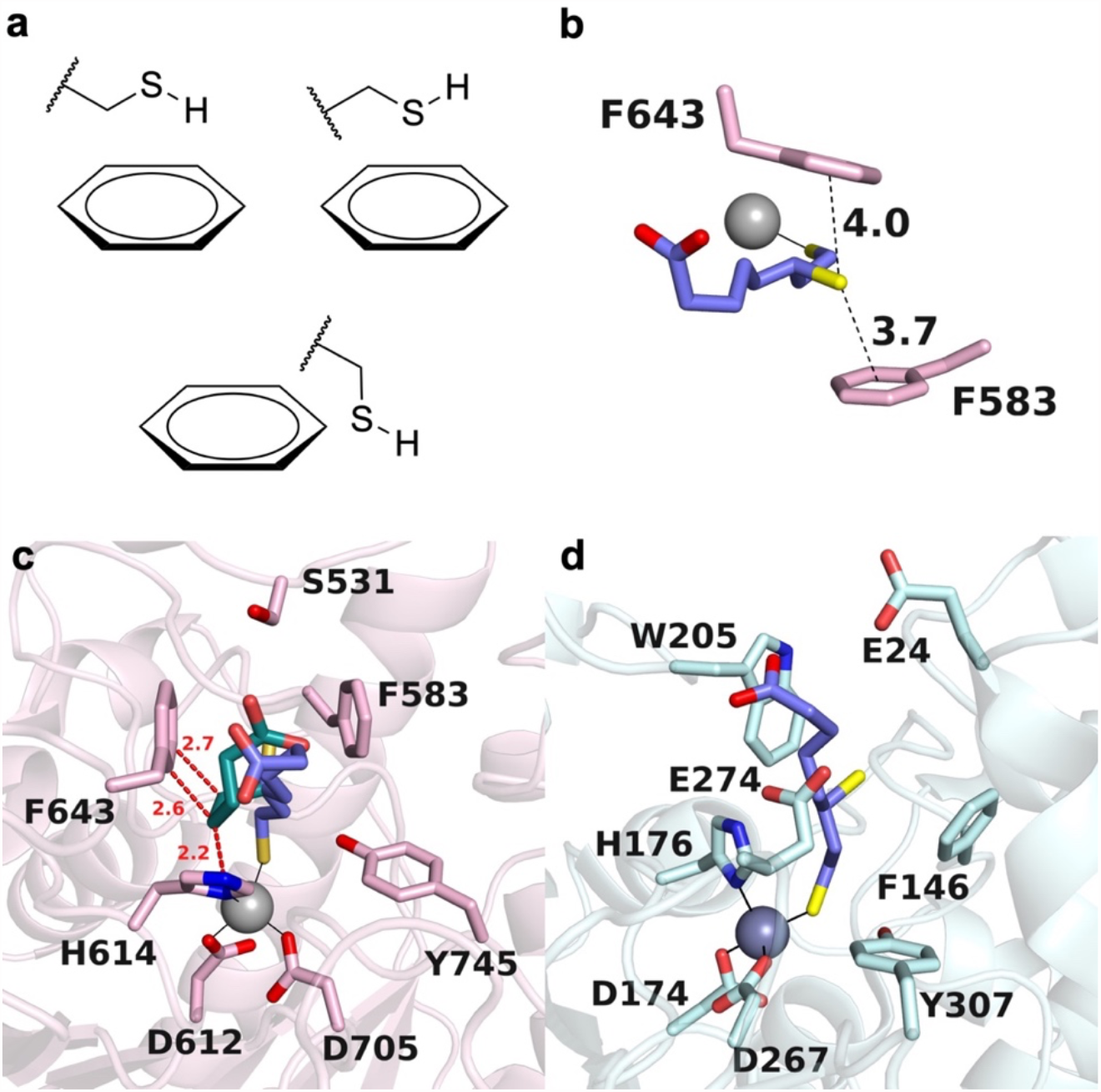
S-π interactions. (a) Three possible orientations for S-H---π interactions. (b) distance of S-π interactions between HDAC6 and the C-6 thiol group of R-α-dihydrolipoic acid in monomer A. The C-6 thiol group is located nearly directly over the centroids of each aromatic ring. (c) Model of (*S*)-dihydrolipoamide (C = turquoise) binding in the HDAC6 active site prepared from the structure of the (*R*)-dihydrolipoamide (C = blue) binding mode in monomer A. Steric clashes with (*S*)-dihydrolipoamide are indicated by red dotted lines and distances. (d) Model of (*R*)-dihydrolipoamide bound in the active site of HDAC10, in which the binding conformation of the inhibitor in HDAC6 monomer A was mapped onto the coordinates of HDAC10.

Notably, molecular recognition of the thiol groups dictates stereoselectivity for the binding of (*R*)-dihydrolipoic acid over the binding of (*S*)-dihydrolipoic acid. If the (*S*)-stereoisomer were to bind with the C8 thiolate coordinated to Zn^2+^ and the C6 thiol bound in the aromatic crevice, our modeling studies suggest that the C5 and C4 methylene groups of (*S*)-dihydrolipoic acid would sterically clash with H614 and F643 (Figure 4c).

Interestingly, Lechner and colleagues show that (*R/S*)-dihydrolipoamide (here, with the carboxylate group amidated to form a neutral carboxamide, such that R = H in Figure 1), but not (*R/S*)-dihydrolipoic acid, is an inhibitor of the class IIb isozyme HDAC10.^17^ Like HDAC6, HDAC10 is enriched in the cytosol,^39^ but unlike HDAC6, HDAC10 is a polyamine deacetylase that exhibits narrow substrate specificity for the hydrolysis of *N*^8^-acetylspermidine.^40^ A negatively charged glutamate residue unique to the HDAC10 active site, E274, confers polyamine substrate specificity by engaging the positively charged secondary ammonium group of *N*^8^-acetylspermidine with water-mediated hydrogen bonds.^41^ If the HDAC6 binding conformation of (*R*)-dihydrolipoic acid is modeled into the active site of HDAC10, the carboxylate group of the inhibitor would reside close to the carboxylate group of E274 and also nearby E24, both of which are clearly repulsive electrostatic interactions (Figure 4d). This explains why neutral (*R*)-dihydrolipoamide is a much better inhibitor of HDAC10, thereby verifying the proposal advanced by Lechner and colleagues for HDAC10 inhibitory potency.^17^

Notably, (*R*)-lipoamide is reported to be a substrate for HDAC11, which catalyzes the hydrolysis of the amide linkage with the tethered lysine side chain.^42^ HDAC11 is a lysine fatty-acid deacylase and can process a variety of fatty acid and cofactor conjugates with lysine residues.^42-44^ Thus, different HDAC isozymes may be involved in lipoic acid function in the cell.

With regard to the biological implications of (*R*)-lipoic acid inhibition of HDAC isozymes, Lechner and colleagues report that both (*R*)-lipoic acid and (*R*)-lipoamide inhibit stress granule formation in cancer cells,^17^ an effect that has also been observed for other HDAC inhibitors such as Trichostatin and Tubastatin.^45^ Moreover, increased HDAC activity is observed following induction of oxidative stress,^46^ and HDAC6 is known to play a critical role in stress response.^47^ Accordingly, we conclude that the structure of the HDAC6–(*R*)-dihydrolipoic acid complex reported herein provides the first visualization of how (*R*)-lipoic acid might regulate HDAC function in the stress response.

### Experimental Procedures

#### Enzyme preparation

HDAC6 catalytic domain 2 from *Danio rerio* (henceforth simply “HDAC6”) was expressed and purified as previously detailed,^48^ with the modification of 0 mM imidazole in Buffer A.

#### Isothermal titration calorimetry

Enthalpograms were measured using a MicroCal iTC 200 isothermal titration calorimeter (Malvern Panalytical, Malvern, UK). For (*R/S*)-lipoic acid, (*S*)-lipoic acid, and (*R*)-lipoic acid, 240 μM inhibitor was incubated for 3 h in size exclusion (SE) buffer [50 mM 4-(2-hydroxyethyl)-1-piperazineethanesulfonic acid (HEPES) (pH = 7.5), 100 mM KCl, 1 mM tris(2-carboxyethyl)phosphine (TCEP), 5% glycerol] and titrated against 30 μM HDAC6 in SE buffer. The (*R/S*)-lipoamide sample was solubilized in DMSO and incubated for 3 h in SE buffer; 240 μM inhibitor was titrated against 30 μM HDAC6 in SE buffer with 5% DMSO. Twenty 2 μL-injections were made over 40 min, with constant stirring in the sample cell (750 r.p.m.) at 25° C. Background was taken under same conditions without HDAC6 (inhibitor to buffer). Integration, reference subtraction, curve fitting, and figure generation were performed using Origin (OriginLab, Northampton, MA).

#### Crystallography

The HDAC6 complex with (*R*)-lipoic acid was crystallized by the sitting-drop vapor diffusion method. Prior to inhibitor incubation, 4 µL of 40 mM (*R*)-lipoic acid was incubated with 0.4 µL of 1.0 M TCEP in 12.7 µL of buffer [50 mM HEPES (pH 7.5), 100 mM KCl, 5% glycerol (v/v), 1 mM TCEP] for 4 hours. Following this, 22.9 µL of 17.5 mg/mL HDAC6 was added for a total volume of 40 µL. Final concentrations of key components were 10 mg/mL HDAC6, 4 mM (*R*)-lipoic acid, and 10 mM TCEP. A 200 nL drop of precipitant solution [0.2 M potassium citrate tribasic and 20% (w/v) polyethylene glycol 3,350] was combined with a 200 nL drop of protein solution and equilibrated against 80 µL of precipitant solution in the well reservoir at 4° C. Plate-like crystals formed in 2 days. Prior to data collection, crystals were flash-cooled in mother liquor supplemented with 20% ethylene glycol.

X-ray diffraction data were collected on the NSLS-II FMX beamline at Brookhaven National Laboratory. Data was indexed using XDS and scaled using Aimless.^49,50^ The initial electron density map was phased by molecular replacement using Phaser^51^ with the atomic coordinates of unliganded HDAC6 (PDB 5EEM)^21^ as a search probe for rotation and translation function calculations (PDB 5EEM). The protein model was adjusted using Coot and refined in Phenix.^52,53^ (*R*)-lipoic acid was fit to the electron density map in the later stage of refinement.

Validation of the final model was performed with MolProbity.^54^ Data collection and refinement statistics are recorded in Table 1.

**Table 1.** Data collection and refinement statistics for the HDAC6–(*R*)-lipoic acid complex.

|  |  |
| --- | --- |
| Space group | $P2_12_12_1$ |
| a,b,c (Å) | 74.89, 92.16, 96.67 |
| $\alpha, \beta, \gamma$ (°) | 90.0, 90.0, 90.0 |
| $R_{\text{merge}}^b$ | 0.218 (0.655) |
| $R_{\text{pim}}^c$ | 0.106 (0.312) |
| $CC_{1/2}^d$ | 0.992 (0.965) |
| Redundancy | 9.2 (9.9) |
| Completeness (%) | 99.6 (100) |
| $I/\sigma$ | 8.7 (4.9) |
| Refinement |  |
| Resolution (Å) | 29.6–2.4 (2.49–2.40) |
| No. unique reflections | 26723 (2782) |
| $R_{\text{work}}/R_{\text{free}}^e$ | 0.207/0.248 (0.232/0.307) |
| Number of atoms<sup>f</sup> |  |
| Protein | 5408 |
| Ligand | 27 |
| Solvent | 132 |
| Average B factors (Å<sup>2</sup>) |  |
| Protein | 18 |
| Ligand | 21 |
| Solvent | 18 |
| RMS deviations |  |
| Bond lengths (Å) | 0.003 |
| Bond angles (°) | 0.56 |
| Ramachandran plot<sup>g</sup> |  |
| Favored | 96.71 |
| Allowed | 3.29 |
| Outliers | 0.00 |

<sup>a</sup>Values in parentheses refer to the highest-resolution shell of data.
<sup>b</sup> $R_{\text{merge}} = \sum_h \sum_i |I_{i,h} - \langle I \rangle_h| / \sum_h \sum_i I_{i,h}$ , where $\langle I \rangle_h$ is the average intensity calculated for reflection $h$ from $i$ replicate measurements.
<sup>c</sup> $R_{\text{p.i.m.}} = (\sum_h (1/(N-1))^{1/2} \sum_i |I_{i,h} - \langle I \rangle_h|) / \sum_h \sum_i I_{i,h}$ , where $N$ is the number of reflections and $\langle I \rangle_h$ is the average intensity calculated for reflection $h$ from replicate measurements.
<sup>d</sup>Pearson correlation coefficient between random half-datasets.
<sup>e</sup> $R_{\text{work}} = \sum ||F_o| - |F_c|| / \sum |F_o|$ for reflections contained in the working set. $|F_o|$ and $|F_c|$ are the observed and calculated structure factor amplitudes, respectively. $R_{\text{free}}$ is calculated using the same expression for reflections contained in the test set held aside during refinement.
<sup>f</sup>Per asymmetric unit.
<sup>g</sup>Calculated with MolProbity.

## Data availability

Atomic coordinates and structure factor amplitudes of the HDAC6–(*R*)-lipoic acid complex have been deposited in the Protein Data Bank (www.rcsb.org) with accession code 8TQ0.

## Acknowledgments

This work is based on research conducted at the FMX beamline of the National Synchrotron Light Source II, a DOE Office of Science User Facility operated for the DOE Office of Science by Brookhaven National Laboratory under Contract DE-SC0012704. The Center for BioMolecular Structure (CBMS) is primarily supported by the National Institutes of Health, NIGMS, through a Center Core P30 Grant (P30GM133893) and by the DOE Office of Biological and Environmental Research (KP1607011).

## Author contributions

P.R.W., conceptualization and X-ray crystallography; J.G.S., isothermal titration calorimetry; P.R.W., J.G.S., and D.W.C., data analysis and manuscript preparation.

## Funding and additional information

This research was supported by NIH grant GM49758. The content is solely the responsibility of the authors and does not necessarily represent the official views of the NIH.

## Abbreviations

The abbreviations used are:

CoA: Coenzyme A
HEPES: 4-(2-hydroxyethyl)-1-piperazineethanesulfonic acid
HDAC6: histone deacetylase 6
HDAC10: histone deacetylase 10
ITC: isothermal titration calorimetry
PDB: Protein Data Bank
TCA cycle: tricarboxylic acid cycle
TCEP: tris(2-carboxyethyl)phosphine

